# Improving Whole-Brain Neural Decoding of fMRI with Domain Adaptation

**DOI:** 10.1101/375030

**Authors:** Shuo Zhou, Christopher R. Cox, Haiping Lu

**Author notes:** Email addresses:* (Shuo Zhou), (Christopher R. Cox), (Haiping Lu).

## Abstract

In neural decoding, there has been a growing interest in *machine learning* on *whole-brain* functional magnetic resonance imaging (fMRI). However, the size discrepancy between the feature space and the training set poses serious challenges. Simply increasing the number of training examples is infeasible and costly. In this paper, we proposed a *domain adaptation framework for whole-brain fMRI* (DawfMRI) to improve whole-brain neural decoding on *target* data leveraging pre-existing *source* data. DawfMRI consists of three steps: 1) *feature extraction* from whole-brain fMRI, 2) source and target *feature adaptation*, and 3) source and target *classifier adaptation*. We evaluated its eight possible variations, including two non-adaptation and six adaptation algorithms, using a collection of seven task-based fMRI datasets (129 unique subjects and 11 cognitive tasks in total) from the OpenNeuro project. The results demonstrated that appropriate source domain can help improve neural decoding accuracy for challenging classification tasks. The best-case improvement is 8.94% (from 78.64% to 87.58%). Moreover, we discovered a plausible relationship between psychological similarity and adaptation effectiveness. Finally, visualizing and interpreting voxel weights showed that the adaptation can provide additional insights into neural decoding.

## 1 Introduction

Functional magnetic resonance imaging (fMRI) is a medical imaging technique that records the Blood Oxygenation Level Dependent (BOLD) signal caused by changes in blood flow (Ogawa et al., 1990). Typically, an fMRI sequence is composed of MRI volumes sampled every few seconds, where each MRI volume has over 100,000 voxels and each voxel represents the aggregate activity within a small volume (2 – 3mm^3^). fMRI can measure the neural activity associated with multiple cognitive behaviors and examine brain functions in healthy relative to disordered individuals. Distinguishing functional brain activities can be framed as classification problems and solved with machine learning techniques, e.g. classifying clinical populations (Norman et al., 2006; Tong & Pratte, 2012) or cognitive tasks (Cheng et al., 2015c; Gheiratmand et al., 2017). However, in such settings, the number of fMRI training examples available for machine learning is relatively small compared to the feature (voxel) space, typically less than one hundred examples per brain state. This size discrepancy makes accurate prediction a challenging problem.

Traditionally, this challenge is dealt with by preselecting voxels that belong to regions of interest (ROIs) based on prior work and established knowledge of domain experts (Poldrack, 2007), or by performing a “searchlight” analysis (Kriegeskorte et al., 2006). While making the problem computationally more tractable, it may ignore a significant portion of information in fMRI, and miss potentially valid and superior solutions in the first place. Recently, studies of *whole-brain fMRI* are becoming increasingly popular (Allen et al., 2014; Gonzalez-Castillo et al., 2015; Vu et al., 2015). This approach not only broadens the scope of potentially important differences between cognitive states, but also lifts the need for a priori assumptions about which parts of the brain are most relevant. Whole-brain fMRI analysis can take all available information into account in a more *data-driven* workflow, however they can be severely hindered by their small training sets.

While any individual fMRI dataset provides only a few training examples, growing public data repositories collectively contain much more training examples, such as OpenNeuro^1^ and the Human Connectome Project^2^. While every neuroimaging experiment is importantly different, many recruit similar sets of cognitive functions. If training examples from related, pre-existing datasets can be leveraged to improve the performance and interpretability of decoding models, it would unlock immense latent power in big data resources that already exist in the neuroimaging community.

*Domain adaptation*, or more broadly *transfer learning*, is an emerging machine learning scheme for solving such a problem (Pan & Yang, 2010). It aims at improving the classification performance in a particular classification problem by utilizing the knowledge learned from different but related data. In domain adaptation terminology, the data to be classified are called the *target domain data*, while the data to be leveraged are called the *source domain data* (Pan & Yang, 2010). The effectiveness of domain adaptation has been shown in varied domains such as computer vision and natural language processing (Patel et al., 2015; Weiss et al., 2016).

Domain adaptation has also been applied to brain imaging data in decoding cognitive states. For example, Zhang et al. (2018) proposed two transfer learning approaches by making use of shared subjects between target and source datasets. They employed two factorization models (Varoquaux et al., 2011; Chen et al., 2015) to learn subject-specific bases, which are assumed to be invariant across datasets. Using a source dataset with shared subjects can help the models learn better subject-specific bases, and therefore improve the prediction accuracy. However, the approaches are not applicable when no subjects are shared between datasets, which is common for multi-site data sharing projects.

There is another related multi-task learning (MTL) approach for neural decoding. Rao et al. (2013) proposed sparse overlapping sets lasso (SOS Lasso) for fMRI, with an MTL approach. MTL is a branch of transfer learning, which aims at improving the performance of all tasks considered and does not differentiate source and target domains. In the context of SOS Lasso, the multiple “tasks” are datasets associated with each of several *participants* of *the same* experiment. The tasks are related as in multitask group lasso (Yuan & Lin, 2006), but groups can be overlapped with one another, and features can be sparsely selected both within and across groups. This technique uses all data relevant to a specific classification problem, but it does not leverage similarities among different classification problems.

In diagnosing brain diseases or disorders, Li et al. (2018) developed a deep transfer learning neural network to improve the autism spectrum disorder classification by leveraging an autoencoder (Vincent et al., 2010) trained on the data of a large number of healthy subjects. Ghafoorian et al. (2017) applied deep learning based domain adaptation to brain MRI for lesion segmentation with a convolutional neural network trained on a source domain of 280 patients and the last few layers fine-tuned on a target domain of 159 patients. Wachinger et al. (2016) proposed an instance re-weighting framework to improve the accuracy of Alzheimer’s Disease (AD) diagnosis by making the source domain data to have similar distributions as target domain data. Cheng et al. (2012, 2015a,b, 2017) proposed several workflows to perform domain adaptation to improve AD diagnosis accuracy by leveraging data of mild cognitive impairment, which is considered as the early stage of AD.

Despite the progresses in the broad domain of neuroimaging, to the best of our knowledge, domain adaptation has not been studied systematically for whole-brain fMRI data. This paper proposes a ***D**omain **a**daptation framework for **w**hole-brain **fMRI*** (DawfMRI) to improve the performance in a *target domain* classification problem with the help of *source domain* data. This enables systematic study of domain adaptation for whole-brain fMRI to evaluate and further develop feasible solutions. It can also help discover novel findings, understand how domain adaptation works in the context of neuroimaging, and identify key technical challenges. Our main contributions are twofold:

1. Methods: We formulated the DawfMRI framework consisting of three steps: 1) *feature extraction* from whole-brain fMRI, 2) source and target *feature adaptation*, and 3) source and target *classifier adaptation*. Under this framework, we identified a state-of-the-art realization of each step and evaluated all eight possible variations, including two non-adaptation and six adaptation algorithms. Our source code is available at: https://github.com/szl44/DawfMRI.
2. Results: We designed experiments systematically using a collection of tasks from the OpenNeuro/OpenfMRI^3^ project. Results demonstrated a promising way of leveraging existing source data to improve neural decoding on target data. We discovered a plausible relationship between psychological similarity and adaptation effectiveness and revealed additional insights obtained via adaptation.

## 2 Materials and Methods

### 2.1 OpenfMRI Data

We chose seven OpenfMRI datasets^4^ used in the study reported in (Poldrack et al., 2013). Table 1 lists the details of the selected datasets. OpenfMRI is a public fMRI data sharing project. It provides whole-brain task-based functional MRI data, as well as structural MRI data and metadata. Both functional and structural images are in the NIfTI format. The metadata record experiment-related information, such as onset time, length, and weighting.

**Table 1:** List of OpenfMRI datasets used in our experiments (ACN denotes accession number, #Sub denotes the number of subjects in each run, and #Run denotes the number of runs used in our experiments).

| Task ID | ACN | #Sub | #Run | Task & Contrast Description |
| --- | --- | --- | --- | --- |
| 01 | ds001 | 16 | 2 | Balloon analog risk task: Parametric pump effect vs. control |
| 02 | ds002 | 17 | 2 | Probabilistic classification: Task vs. baseline |
| 03 | ds002 | 17 | 2 | Deterministic classification: Feedback vs. baseline |
| 04 | ds002 | 17 | 2 | Mixed event-related probe: Task vs. baseline |
| 05 | ds003 | 13 | 1 | Rhyme judgement: Task vs. baseline |
| 06 | ds005 | 16 | 2 | Mixed-gambles task: Parametric gain response |
| 08 | ds007 | 20 | 2 | Stop signal task: Letter classification vs. baseline |
| 09 | ds007 | 19 | 2 | Stop signal task: Letter naming vs. baseline |
| 10 | ds007 | 20<sup>5</sup> | 2 | Stop signal task: Pseudoword naming vs. baseline |
| 21 | ds101 | 21 | 2 | Simon task: Incorrect vs. correct |
| 22 | ds102 | 26 | 2 | Flanker task: Incongruent vs. congruent |

The seven datasets contain 11 tasks in total. We used the same task ID as Poldrack et al. (2013). Tasks 2, 3, 4 and tasks 8, 9, 10, respectively, are contributed by the same subjects. We used the original version (revision version 1.0.0) of each dataset. In total, there are 202 fMRI sequences from 129 unique subjects for run 1 and 188 sequences from 116 unique subjects for run 2.

### 2.2. Data Preprocessing Pipeline

To process the data from OpenfMRI, we implemented a standard preprocessing pipeline using FSL (Jenkinson et al., 2012) based on the processing stream^6^ implemented by Poldrack et al. (2013). The output of one step will be the input of next step. As shown in Table 2, the pipeline has five steps:

1. Perform motion correction on the BOLD signal sequences from OpenfMRI using MCFLIRT (FSL).
2. Perform brain extraction using BET (FSL).
3. Perform first-level analysis to generate statistical parametric maps (SPMs) (Friston et al., 1994, 1998) of contrasts for each experiment condition using FEAT (FSL). FSL design files are generated from the OpenfMRI onsets files using the custom code.
4. Align the spatially normalized Z statistic maps obtained in Step 3 with the MNI152 standard image using featregapply (FSL). The data dimension is standardized to 109 × 109 × 91 (2*mm*^3^) voxels.
5. Vectorize the voxels from the Z statistic maps that fall within the standard MNI152 T1 2*mm* brain mask (distributed with FSL) using the Python package Nibabel (Brett et al., 2017).

**Table 2:**
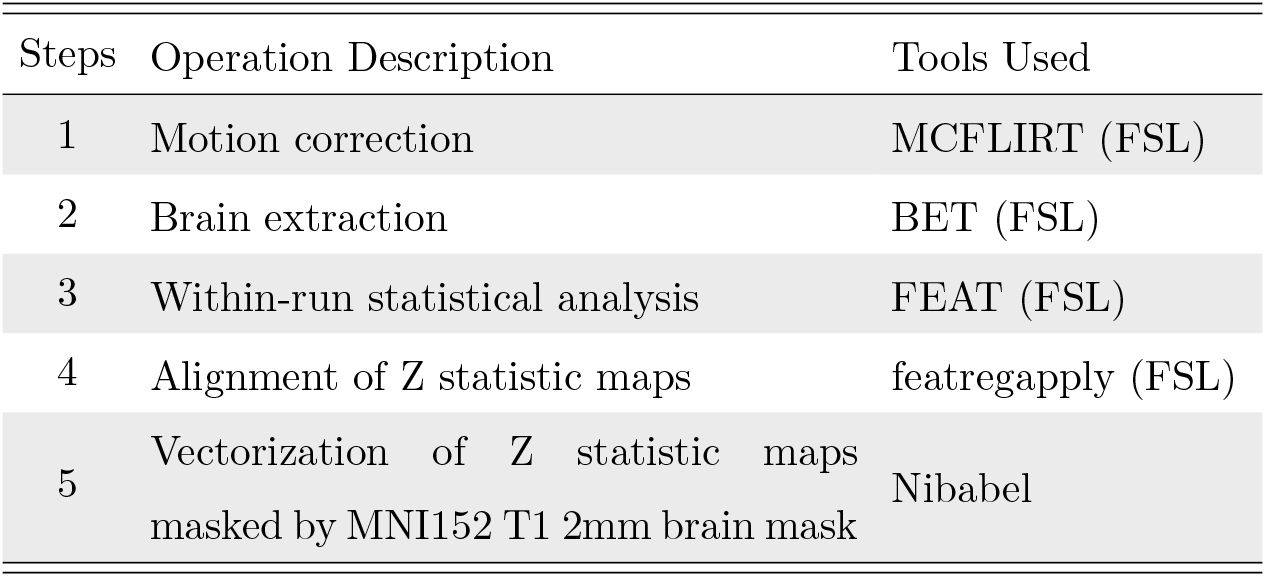
Preprocessing pipeline for the selected OpenfMRI data.

| Steps | Operation Description | Tools Used |
| --- | --- | --- |
| 1 | Motion correction | MCFLIRT (FSL) |
| 2 | Brain extraction | BET (FSL) |
| 3 | Within-run statistical analysis | FEAT (FSL) |
| 4 | Alignment of Z statistic maps | featregapply (FSL) |
| 5 | Vectorization of Z statistic maps<br>masked by MNI152 T1 2mm brain mask | Nibabel |

The contrasts associated with each task represent differences between the primary tasks and some baseline conditions. Rather than considering the influence of various baseline conditions, we conducted our experiments using the same single contrast per task as Poldrack et al. (2013). The contrasts used are also reported in Table 1.

### 2.3 Domain Adaptation for Whole-Brain fMRI

We propose a domain adaptation framework for whole-brain fMRI (DawfMRI) as shown in Fig. 1. This framework consists of three steps: feature extraction, feature adaptation, and classifier adaptation.

**Figure 1:**
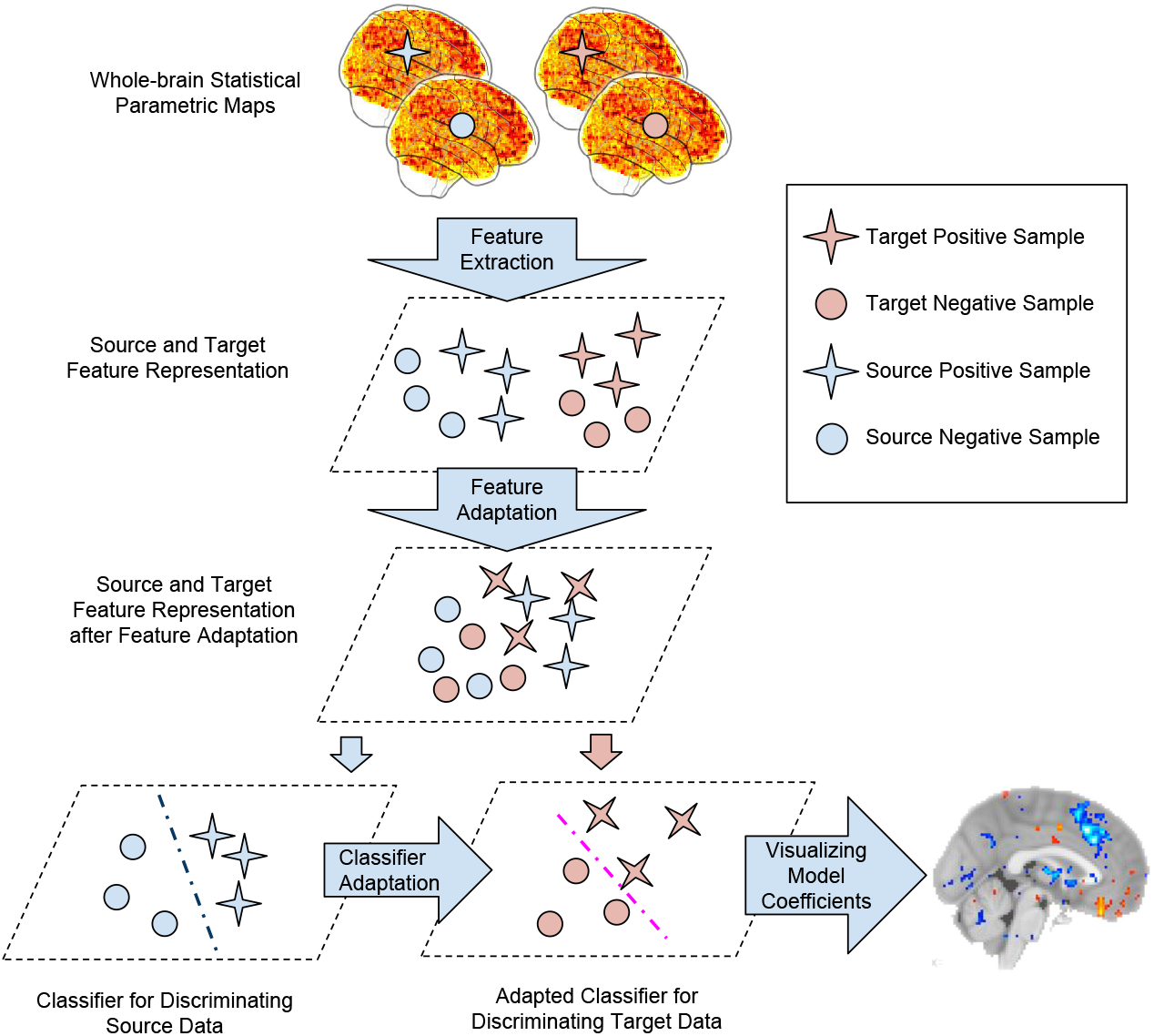
The proposed domain adaptation framework for whole-brain fMRI consists of three steps: feature extraction, feature adaptation, and classifier adaptation, e.g. via independent component analysis, transfer component analysis, and cross-domain SVM, respectively. Learned model can be visualized on a brain atlas for cognitive interpretation.

- *Feature extraction* aims to distill informative and non-redundant features from high-dimensional data to accelerate computation, reduce overfitting, and facilitate interpretation. Whole-brain fMRI is very high-dimensional. However, there are not as many meaningful components as the number of voxels. Thus there is high redundancy and feature extraction can be applied to reduce the dimension by identifying a more compact set of informative features, e.g. with principal component analysis or independent component analysis.
- *Feature adaptation* is a domain adaptation scheme that utilizes the source domain samples for target model training. The motivation is to leverage the samples from a related (source) domain when the information provided by the target domain samples is limited for training a good model. However, a classifier trained on source domain data will typically perform poorly on the target domain classification. This is due to domain feature distribution mismatch, which means that the features in target and source domains do not follow the same probability distribution. The objective of feature adaptation is to minimize this mismatch by feature mapping or reweighting (Pan & Yang, 2010). After performing feature adaptation, the adapted samples from source domain can be used as additional samples for training the target model.
- *Classifier adaptation* is another domain adaptation scheme that aims at improving the classifier performance in a target domain using the knowledge, such as coefficients or parameters, from a pre-trained classifier. The motivation is that when the information provided by target domain data is limited to train a good classifier, the discriminative information that a classifier learned from source data can be leveraged to train a better target classifier.

There is a key difference between feature adaptation and classifier adaptation. The goal of feature adaptation is to make the source and target domain data similar. Classifier adaptation, in contrast, involves fitting a model to the source domain data, and using this model to set priors for another model of the target domain. When feature adaptation is used without subsequent classifier adaptation, the feature-adapted source domain is used directly as if it contains additional training examples in the feature-adapted target domain. That is, a single model will be fit to the feature-adapted source and target domain data.

Each step of DawfMRI can be optional. If all the three steps are skipped, we train a classifier directly on the whole-brain data. To study DawfMRI systematically, we employ a state-of-the-art method for each step in DawfMRI: independent component analysis (ICA) (Comon, 1994) for feature extraction, transfer component analysis (TCA) (Pan et al., 2011) for feature adaptation, and cross-domain SVM (CDSVM) (Jiang et al., 2008) for classifier adaptation. Table 3 lists the key notations used for easy reference in the following presentation of these methods.

**Table 3:** Notations and descriptions.

| Notation | Description |
| --- | --- |
| $D$ | Input feature dimension |
| $\mathbf{D}_l^t$ | Labeled target data |
| $d$ | Output (lower) feature dimension |
| $\mathbf{H}$ | Centering matrix |
| $\mathbf{I}$ | Identity matrix |
| $\mathbf{K}$ | Kernel matrix |
| $k(\cdot, \cdot)$ | Kernel function, e.g. linear, Gaussian, polynomial |
| $n_T, n_S$ | Target/source sample size |
| $\mathbf{U}$ | Feature mapping/transformation matrix |
| $\mathbf{V}^s$ | Source support vectors |
| $\mathbf{w}$ | Classifier coefficients |
| $\mathbf{X}_T, \mathbf{X}_S$ | Target/source data |
| $\mathbf{Z}_T, \mathbf{Z}_S$ | Learned target/source representation |

#### 2.3.1 Feature Extraction by ICA

ICA is a popular method for fMRI feature extraction. We followed the procedure of performing ICA in (Poldrack et al., 2013) in our experiments. Spatial smoothing was performed on the SPMs obtained from the pipeline in Sec. 2.2. Then ICA was performed on smoothed whole-brain SPMs using MELODIC (Beckmann & Smith, 2004), the ICA tool in FSL.

We performed ICA on data from all seven datasets for better estimation quality. The objective of this step is to extract informative low-dimensional features from high-dimensional input rather than adaptation. It is unaware of the domain differences and does not attempt to utilize this information. Therefore, there is no adaptation in this step and all data share a common ICA feature space.

#### 2.3.2 Feature Adaptation by TCA

TCA aims to find a feature mapping to minimize the mismatch between target and source distributions, using the Maximum mean discrepancy (MMD) (Borgwardt et al., 2006) as the distribution mismatch metric. Given source domain data **X**_*S*_ ∈ ℝ^*D*×*n_s_*^, target domain data **X**_*T*_ ∈ ℝ^*D*×*n_T_*^, where *n_S_* and *n_T_* denote the number of samples of **X**_*S*_ and **X**_*T*_ respectively, and *D* denotes the input feature dimension, MMD between the two domains is

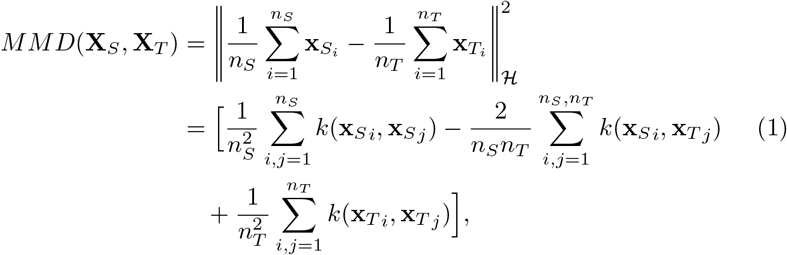

where 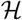 denotes a reproducing kernel Hilbert space (RKHS) (Berlinet & Thomas-Agnan, 2011), *k*(·, ·) denotes a kernel function, such as linear, Gaussian, and polynomial, **x**_*Si*_ and **x**_*T_j_*_ are the *i*th and *j*th sample of **X**_*S*_ and **X**_*T*_, respectively. TCA assumes that the domain difference is caused by marginal distribution mismatch, i.e, *P*(**X**_*S*_) ≠ *P*(**X**_*T*_). Hence, the objective is to learn new feature representations **Z**_*S*_ and **Z**_*T*_ by mapping the input data to a feature space where the MMD between the two domains is minimized, i.e., *P*(**Z**_*S*_) ≈ *P*(**Z**_*T*_). Equation (1) can be rewritten as *MMD*(**X**_*S*_, **X**_*T*_) = *tr*(**KL**), where

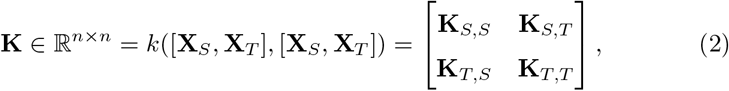

**L** ∈ ℝ^*n* × *n*^ is:

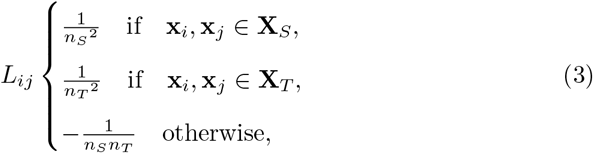

and *n* = *n_S_* + *n_T_*. To minimize MMD, TCA employs a dimension reduction approach. A matrix Ũ ∈ ℝ^*n*×*d*^ is used to map the kernel features to a *d*-dimensional space (*d* ≪ *n_S_* + *n_T_*), which results in a new kernel matrix

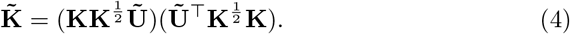

We can consider 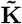 as the inner product of the new representation **Z** = [**Z**_*S*_, **Z**_*T*_] ∈ ℝ^*d*×*n*^. Then 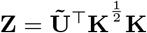. Let 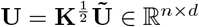, we obtain

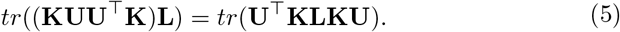

In addition, covariance matrix **U**^⊤^**KHKU** (Fukunaga, 1990) is employed for preserving the properties (variance) of **X**_*S*_ and **X**_*T*_, where **H** ∈ ℝ^*n*×*n*^ is a centering matrix (Marden, 2014). Consequently, the learning objective becomes

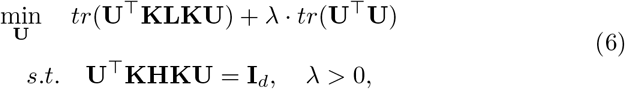

where **I**_*d*_ ∈ ℝ^*d*×*d*^ is an identity matrix, *λ* is a tradeoff parameter for regularization. Denoting Γ = *diag*(*γ*_1_,…, *γ_d_*) as Lagrange multipliers, we can derive the Lagrange function for Eq. (6) as

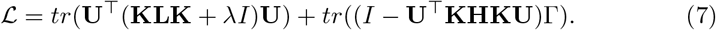

Setting 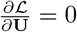, we obtain the following generalized eigendecomposition problem

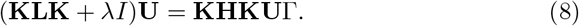

Finally, **U** can be learned by solving Eq. (8) for the *d* smallest eigenvectors.

#### 2.3.3. Classifier Adaptation by CDSVM

CDSVM is an SVM classifier that utilizes the source support vectors learned by a standard SVM on source domain samples to find a better decision boundary for target samples. It re-weights each source support vector according to its average distance to the target (training) feature vectors, and then the target classifier will be trained with the target training samples and re-weighted source support vectors. We learn the decision function of CDSVM *f*(**x**) = **w**^⊤^**x** by optimizing the following objective:

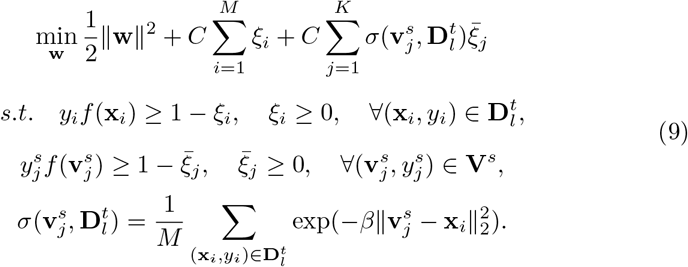

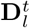 represents the labeled (training) target domain data. **V**^*s*^ denotes a matrix composed of all source support vectors. *M* is the number of samples in 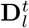. K is the number of source support vectors in **V**^*s*^. 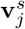 is the *j*th source support vector in **V**^*s*^. *ξ* and 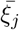 are slack variables for the *i*th target feature vector and *j*th source support vector respectively. *C* is a hyperparameter controlling the tradeoff between the slack variable penalty and the SVM soft margin. 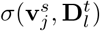 is a function that evaluates the distance between 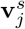 and 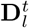. *β* is a hyperparameter controlling the influence of the source support vectors. Larger value of *β* leads to less influence of source support vectors, and vice versa.

### 2.4. Visualizing Model Coefficients for Interpretation

The final classifier coefficients (or weights) indicate the significance of the corresponding features for a classification problem. Visualizing them in the brain voxel space can help us gain some insights into which areas contribute more to prediction performance. To achieve this, we chose a linear kernel in TCA, SVM and CDSVM and developed a method to map the classifier coefficients back to voxel weights in the original brain voxel space for interpretation, as shown in Fig. 2.

**Figure 2:**
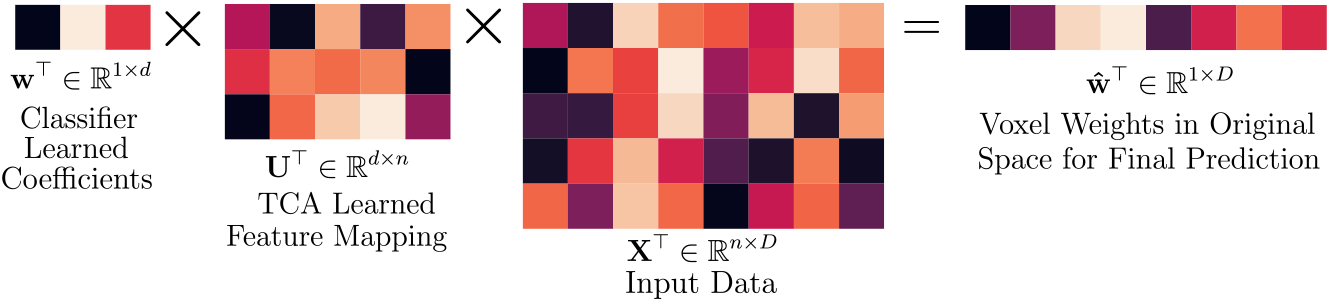
Mapping classifier coefficients **w** ∈ ℝ^*d*×1^ back to voxel weights 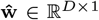 in the voxel feature space.

In the following, we detail how to map the model coefficients of TCA+SVM (or TCA+CDSVM) to the original voxel space. If feature extraction is performed, e.g. in the case of ICA+TCA+SVM, one more step will be needed to map the coefficients in the ICA space to the voxel space, e.g. by GIFT (Calhoun et al., 2001).

Using the same notations above, we have **X** ∈ ℝ^*D*×*n*^ composed of target and source data, TCA has learned a feature transformation **U** ∈ ℝ^*n*×*d*^, and SVM has learned coefficients **w** ∈ ℝ^*d*×1^. According to Eq. (5), the TCA features can be represented as **Z** = **U**^⊤^**X**^⊤^**X** ∈ ℝ^*d*×*n*^. For a sample x ∈ ℝ^*D*×1^, its transformed feature **z** = **U**^⊤^**X**^⊤^x ⊤ ℝ^*d*×1^. The predicted class is

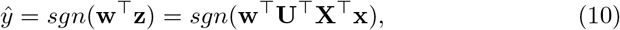

where *sgn* is the sign function (1 for positive values, −1 for negative values). Let 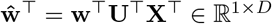, we obtain

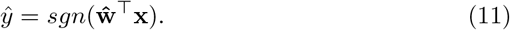

Hence, 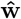 contains the weights corresponding to each voxel in the whole brain space for final prediction.

## 3 Experimental Results

### 3.1. Experiment Settings and Evaluation Methods

Our experiments will focus on using domain adaptation to improve performance on challenging binary classification problems that require distinguishing brain states associated with different cognitive tasks. Thus, we only considered the most basic scenario, specifically, both target and source classification problems are binary. We studied DawfMRI in the setting of one-to-one domain adaptation only. This means that one target domain will be supplemented by only one set of source domain data.

#### 3.1.1. Algorithm Setup

We evaluated eight possible variations of the DawfMRI framework, including two non-adaptation algorithms: 1) SVM and 2) ICA+SVM, and six adaptation algorithms: 3) CDSVM,4) ICA+CDSVM, 5) TCA+SVM, 6) TCA+CDSVM, 7) ICA+TCA+SVM, and 8) ICA+TCA+CDSVM.

For feature adaptation only algorithms (5 & 7), i.e., performing TCA without using CDSVM, an SVM was fit to both of the feature-adapted source and target domain data. For algorithms using CDSVM (3, 4, 6, and 8), we trained an SVM on the source domain, and then used the learned source support vectors as additional input knowledge for training CDSVM on the target domain.

As mentioned in Sec. 2.4, we chose a linear kernel in TCA, SVM, and CDSVM for easy interpretation. We optimized hyperparameters on regular grids of log scale for each algorithm, with a step size of one in exponent. We searched for the best *C* and *μ* values within the range [10^-3^,10^3^] and [10^-5^,10^5^] for SVM and TCA, respectively. For CDSVM, we grid-searched for the best combination of *C* and *β* values in [10^-3^,10^3^]. We also varied the feature dimension of ICA and TCA output from 2 to 100 (2, 10, 20, 50 and 100), and optimize the relevant algorithms with the best feature dimensions.

#### 3.1.2. Target and Source Domain Setup

Each task (listed in Table 1) is associated with data collected from a number of participants over one or two scanner runs. An SPM expressing a particular contrast between task and baseline exists for each run for each subject. These SPMs comprise the set of possible training examples, and the tasks serve as the category labels. Each pair of tasks is a domain, and our problem is binary classification between two tasks. Each domain can be used as a target (the primary problem that we want to improve performance on) or a source (the secondary problem from which we want to leverage knowledge to help better solve the primary problem). We anticipate the classification problems with lower prediction accuracy to have higher potential of improvement via domain adaptation. Therefore, we selected three most challenging domains, with highly confusable pairs of tasks:

1. Tasks 1 (32 samples) & 22 (56 samples),
2. Tasks 3 (34 samples) & 6 (32 samples),
3. Tasks 6 (32 samples) & 22 (56 samples).

These domains were identified by performing multi-class classification. Specifically, we used a linear SVM to classify all eleven classes of the whole-brain SPMs. Figure 3 shows the 10-fold cross-validation results as a confusion matrix, where an entry (*i, j*) is the number of observations actually in task *i*, but predicted to be in task *j*. This allows us to identify the most confusable pairs. The results indicate that Tasks 1 and 6 were often confused with other tasks. Task 1 was misclassified as Task 22 more than half the time. Task 6 was the least accurately classified overall, of which the samples were often misclassified as Tasks 3, 21, or 22. We noted that samples from Task 3 can also be misclassified as Task 6. We then identified the three aforementioned pairs of tasks as target domains to focus on. These selected pairs are confirmed later to be those benefiting the most from domain adaptation in our adaptation effectiveness study (Sec. 3.2.3).

**Figure 3:**
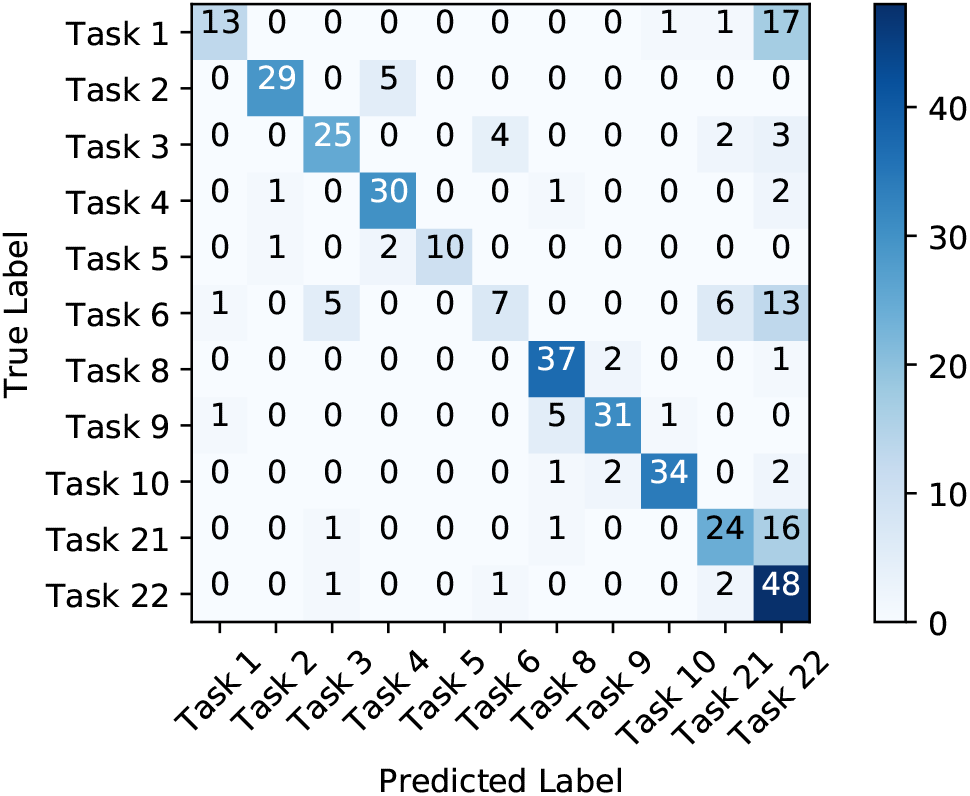
Multi-class classification confusion matrix for linear SVM performance on the whole-brain SPMs. Entry (*i, j*) in the confusion matrix is the number of observations actually in task *i*, but predicted to be in task *j*. Three most challenging pairs of classification tasks (3 vs 6, 6 vs 22, and 1 vs 22) were selected as target domains to perform domain adaptation.

##### Source selection and class labels

When using one pair of tasks as the target domain, the remaining nine tasks are combined pairwise to give 36 unique pairs, each of which is a candidate source domain. For each pair, one task is labeled 1 and the remaining task is labeled −1, also called positive and negative classes, respectively. There are two ways to match source domain labels with target domains labels, i.e., 1 with 1 and −1 with −1, or 1 with −1 and −1 with 1. We studied both cases.

#### 3.1.3. Evaluation Methods

We performed 10 × *k*-fold cross-validation (CV) evaluation, with *k* = 2, 5, and 10, corresponding to using 50%, 80%, and 90% of available target domain samples for training. All training samples were sampled uniformly at random for cross validation. CV was only applied to target domain samples. Source domain samples were all used for training when performing domain adaptation. We will report the mean classification accuracy with standard derivations for performance evaluation. To study the statistical significance of the results obtained by adaptation algorithms compared to those by non-adaptation algorithms, we will report the *p*-values of paired *t*-tests.

Applying CDSVM directly to the whole-brain SPMs ran out of 50GB memory, so we were unable to obtain the results. Experimental results for the *remaining seven algorithms* will be reported in the following section.

### 3.2. Classification Performance

#### 3.2.1. Results on Different Target Domains

Figure 4 shows the 10-CV classification results across the three different target domains. For adaptation algorithms, we tested all possible source domains to report the best results in Fig. 4, with the corresponding sources indicated in the bars. The best results on the three target domain were obtained by TCA+CDSVM, ICA+TCA+SVM, and TCA+CDSVM respectively. The three algorithms outperformed the non-adaptation algorithms significantly (maximum *p*-value < 0.0001 in paired *t*-test). The largest accuracy improvement was obtained by TCA+CDSVM with source 22 vs 9 on the target domain 3 vs 6.

**Figure 4:**
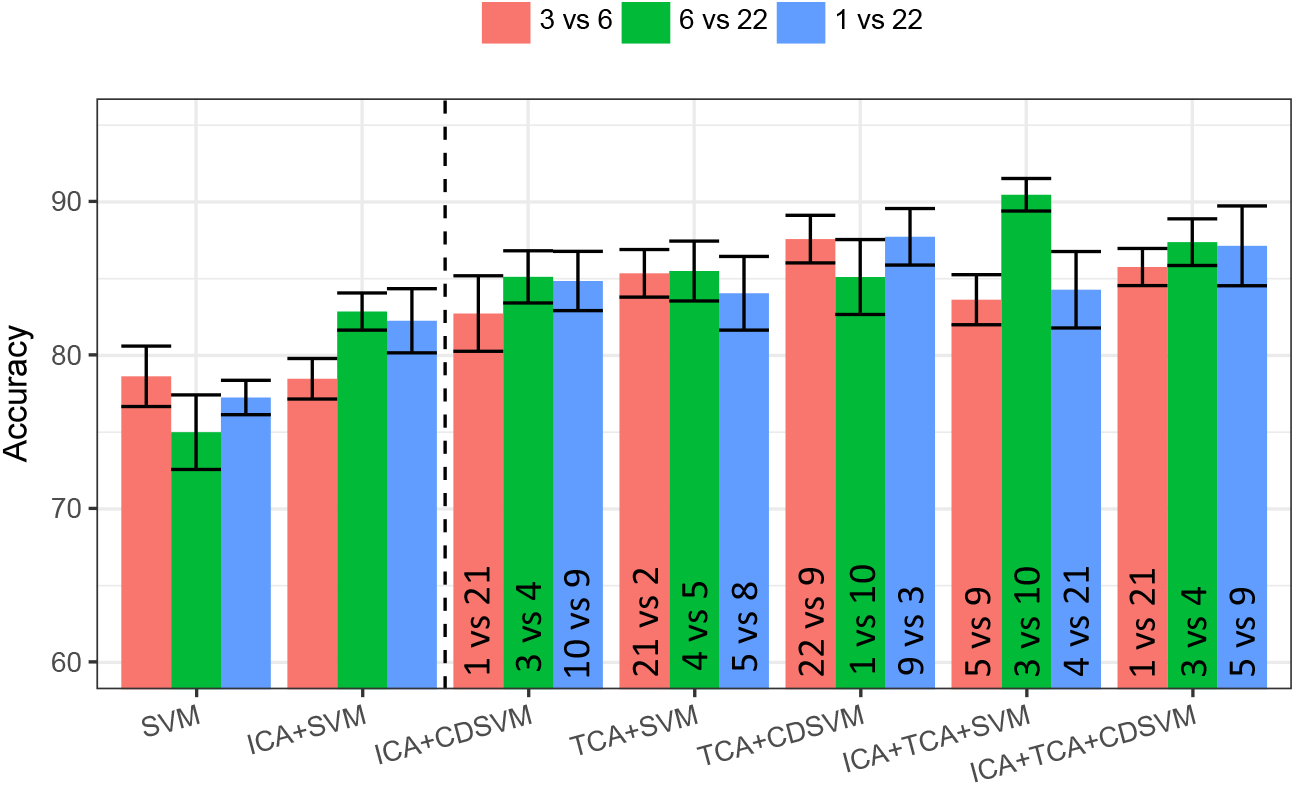
Classification accuracy (in %) of seven algorithms on the three target domains. Non-adaptation (left) and adaptation (right) algorithms are separated by the vertical dashed line. Adaptation algorithms use the best source domains, as indicated in the bars. Error bars indicate the standard derivations.

This improves over the best non-adaptation method (SVM) by **8.94**% (**from 78.64% to 87.58%**).

#### 3.2.2. Effect of Training Sample Size

Next, we fixed the target and source domains to observe how the performance varies with different sizes of training data, specifically, 10-fold, 5-fold, and 2-fold cross-validation. The target domain was fixed to 3 vs 6, and the source domain was fixed to 10 vs 1. Thus, this source domain may not be optimal for adaptation algorithms.

Figure 5 depicts the classification accuracy of seven algorithms for different training samples sizes. TCA+CDSVM achieves the best results for 10-CV and 5-CV, and further paired *t*-test results indicate the improvements over the best non-adaptation algorithms were statistically significant (*p*-value < 0.0001). However, for 2-CV, the accuracy improvement for TCA+SVM (77.37%) and ICA+TCA+CDSVM (77.27%) over ICA+SVM (76.36%) were not statistically significant, with corresponding *p*-values of 0.66 and 0.62 respectively. This indicats that very small training sample size is still challenging, even with adaptation.

**Figure 5:**
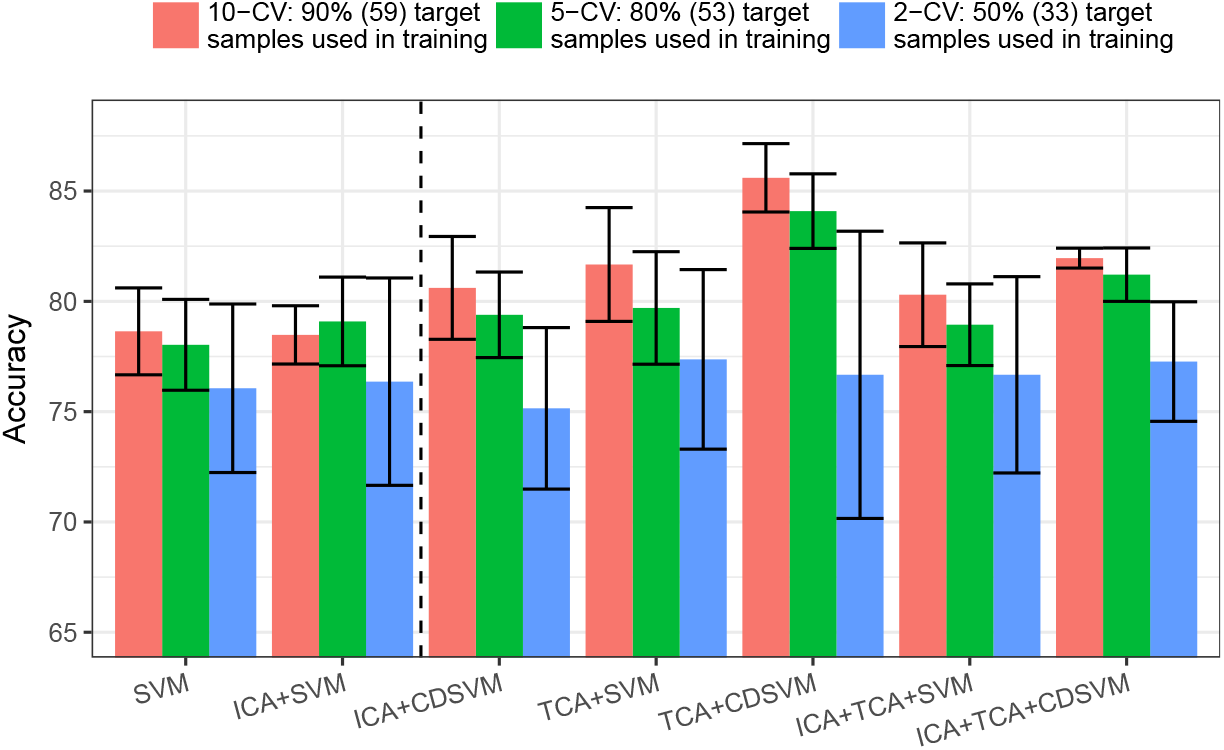
Classification accuracy (in %) of seven algorithms for three cross validation settings on target classification problem task 3 vs task 6 (positive vs negative), with source task 10 vs task 1. Error bars indicate the standard derivations. Non-adaptation (left) and adaptation (right) algorithms are separated by the vertical dashed line.

#### 3.2.3. Sensitivity to Source Domain

For studying the *adaptation effectiveness* of different source domains over different target domains, we applied TCA+CDSVM and SVM to all possible combinations of the 11 tasks as target domains. Figure 6 summarizes the obtained accuracy with the target domains sorted by the largest improvements of TCA+CDSVM over SVM. A psychological interpretation will be given in Sec. 4.1.

**Figure 6:**
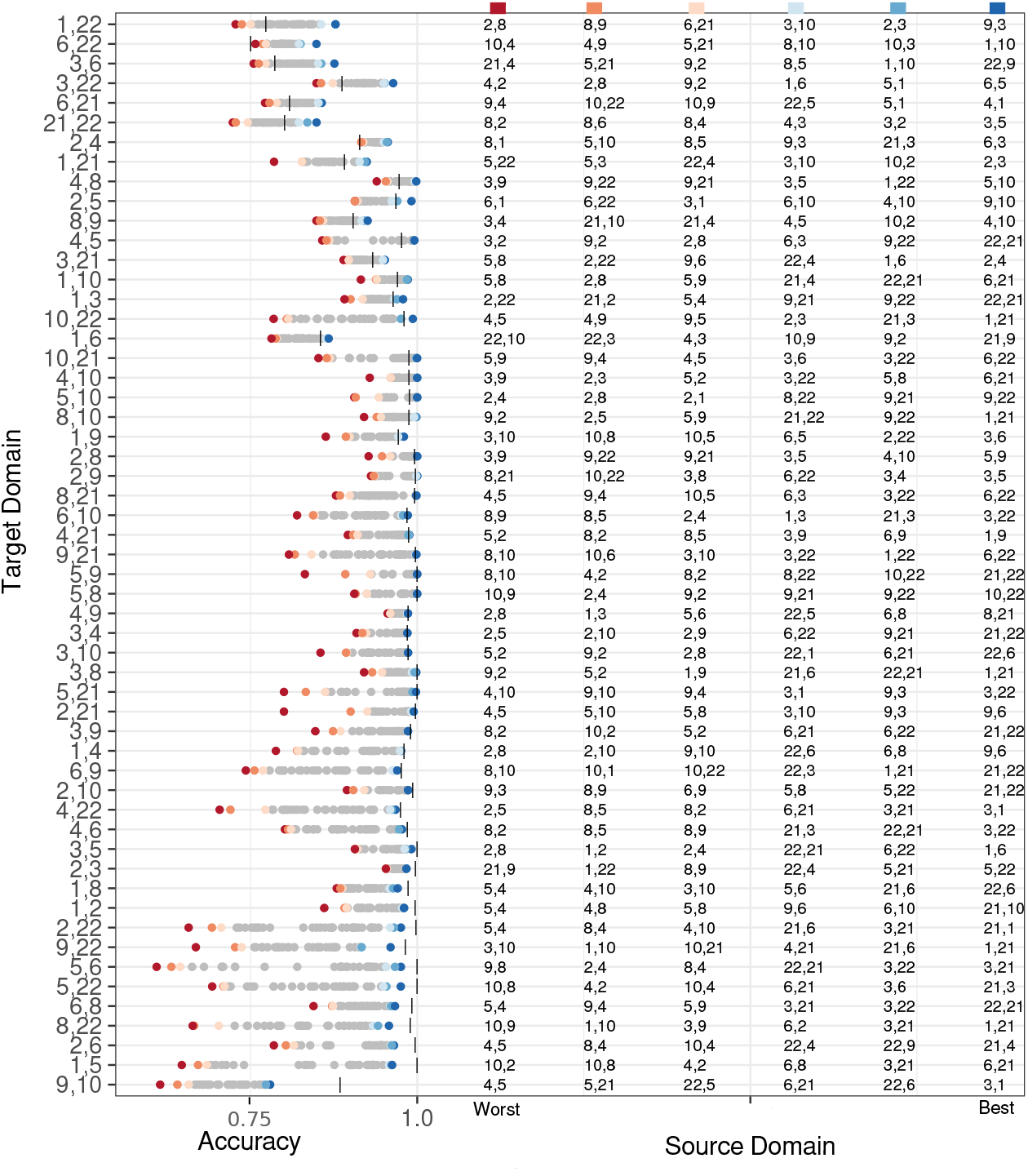
Adaptation effectiveness of TCA+CDSVM (colored dots) over SVM (black vertical bars) across all target domains, with 10-fold cross-validation. Target domains are sorted with respect to the maximum improvement. The top three and bottom three source domains are listed on the right half, in the order of the worst, the second worst, the third worst, the third best, the second best, and the best, from left to right.

Figure 6 shows that the effectiveness of domain adaptation was significantly affected by the source domain, and TCA+CDSVM did not consistently outperform SVM. This is called “negative transfer” (Pan & Yang, 2010).

The figure also shows a negative correlation between the baseline SVM accuracy and the maximum improvements. For higher SVM accuracy, TCA+CDSVM can hardly perform better, instead, it performed worse in many cases (e.g. the target domain of 8 vs 22 and 2 vs 6). This is consistent with our expectation, that domain adaptation is effective for more challenging problems, which motivated our target domain selection strategy in Sec. 3.1.2.

#### 3.2.4. Effectiveness of Each DawfMRI Step

In feature extraction, ICA can extract informative features from high-dimensional whole-brain data. By comparing the results in Fig. 4, we can observe that ICA+SVM outperforms SVM for the targets 1 vs 22 and 6 vs 22. For target 3 vs 6, the accuracy obtained by ICA+SVM is slightly lower than the one obtained by SVM. Considering the much lower dimension of independent components compared with the dimension of whole-brain data, ICA is an effective feature extractor.

In feature adaptation, TCA can also extract features with lower dimension from whole-brain SPMs without performing ICA. Moreover, features extracted by TCA are common and useful across source and target domains. As shown in Figures 4 and 5, both TCA+SVM and ICA+TCA+SVM outperformed non-adaptation algorithms with appropriate sources. This indicates that by performing TCA, samples from appropriate source domains can be used as additional training data for target domain.

In classifier adaptation, CDSVM can improve the accuracy when combined with ICA, TCA, or both. By comparing the results in Figs. 4 and 5, ICA+CDSVM did improve over ICA+SVM consistently, though the amount of improvement was not as large. By contrast, (ICA+)TCA+CDSVM did not outperform (ICA+)TCA+SVM consistently. This indicates that the effectiveness of classifier adaptation tends to be data-dependent.

## 4. Discussion

This section will further analyze DawfMRI with two objectives to facilitate further discussion: 1) exploring whether adaptation effectiveness is related to psychological similarities between tasks, 2) understanding how domain adaptation improves neural decoding by visualizing the model coefficients.

### 4.1. Psychological Interpretation of Source Domain Effectiveness

Domain adaptation effectiveness is closely related to meaningful relationships between the target and source domains. On the other hand, psychological experiments are intrinsically related by the cognitive mechanisms that support the ability to perform the tasks. Hence, we expect the cognitive similarity between a set of tasks to be predictive of whether or not domain adaptation will be effective.

#### 4.1.1. Psychological Similarity Study

We explored the potential relationship between psychological similarity and adaptation effectiveness by modeling the probability of domain adaptation with TCA+CDSVM improving prediction accuracy as a function of the psychological similarity between tasks within and across domains.

To estimate the psychological similarity, we associated each task with a set of cognitive functions that it relies on. The associations were defined by referring to the cognitive concepts in Cognitive Atlas^7^ (Poldrack et al., 2011). Out of the 11 tasks in Table 1, 10 were associated with a number of cognitive functions (min = 5, max = 13, median = 7). The mixed event related probe (task 4) had an incomplete entry so it was excluded from these and further studies. There were 42 functions in total, and each task was represented as a binary feature vector. The psychological similarity between each pair of tasks was computed as the cosine similarity between their feature vectors.

Because each domain is composed of a pair of tasks, the similarity between the target and source domains in each model is associated with four pairwise task similarities. The overall psychological similarity between each pair of domains was estimated by averaging these four pairwise similarities. We will refer to this estimate as the *Cross-Domain Similarity* (CDSim). Moreover, the similarity between the two tasks of the target domain is denoted as the *Target-Domain Similarity* (TDSim), and the similarity between the two tasks of the source domain is denoted as the *Source-Domain Similarity* (SDSim).

This study focused on 504 target and source combinations of the classification problems with no more than 90% in accuracy obtained by whole-brain SVM. Of these 504 models, 261 was improved. We labeled them as ‘improved’ or ‘not improved’ to train a logistic model. This binary outcome was regressed on four variables, CDSim, TDSim, SDSim, and TDSVM_Acc (the accuracy of the standard SVM in the target domain). The variables were standardized to have a mean of zero and standard deviation of one before fitting the model. Table 4 lists the statistics of the four variables.

**Table 4:** Statistics of the four psychological similarity features used for logistic model training, which are target domain similarity (TDSim), source domain similarity (SDSim), cross-domain similarity (CDSim) and target domain SVM accuracy (TDSVM_Acc).

|  | CDSim | TDSim | SDSim | TDSVM_Acc |
| --- | --- | --- | --- | --- |
| Min | -1.78 | -1.02 | -1.21 | -1.60 |
| Median | -0.05 | -0.59 | -0.36 | -0.026 |
| Mean | 0.00 | 0.00 | 0.00 | 0.00 |
| Max | 3.47 | 1.93 | 2.73 | 1.33 |

Table 5 reports the learning outcome of the logistic model, where increasing CDSim increased the probability of improved accuracy. This relationship between psychological similarity and adaptation effectiveness is important. On the one hand, it is consistent with how these adaptation methods are meant to work. On the other hand, it suggests that it may be possible to predict the adaptation effectiveness in advance, without resorting to a post-hoc selection of the source domain through trial and error.

**Table 5:** Logistic model learning for studying the relationship between psychological similarity and adaptation effectiveness. The model was regressed on four variables, TDSim, SDSim, CDSim, and TDSVM_Acc, for predicting improved or not improved.

|  | Beta | Standard Error | *t*-value | *p*-value |
| --- | --- | --- | --- | --- |
| (intercept) | -0.085 | 0.097 | -0.88 | 0.38 |
| TDSim | -0.10 | 0.11 | -0.98 | 0.32 |
| SDSim | -0.20 | 0.11 | -1.89 | 0.057 |
| CDSim | 0.26 | 0.11 | 2.38 | 0.017 |
| TDSVM_Acc | -0.80 | 0.10 | -7.75 | 9.15e-15 |

#### 4.1.2. Source Selection Validation

We also adopted a leave-one-target-domain-out strategy to learn to “select” an appropriate source domain. For the hold-out target domain, we selected the source domain with the highest likelihood of accuracy improvement given by the logistic model. Then we compared the real improvement of using the selected source data against random source selection, i.e., the mean improvement, for TCA+SVM, TCA+CDSVM, and ICA+TCA+SVM. Results showed that psychological similarity based source selection led to 0.0068, 0.0065, and 0.0371 higher classification accuracy than random selection, respectively. Therefore, it can help source selection.

In addition, we did the same analysis to TCA+SVM to compare MMD-based source selection with random source selection. We computed the MMDs for all possible target and source combinations using Eq. (5) after performing TCA, as well as the accuracy of respective TCA+SVM. Results showed that on average, selecting source domains with the smallest MMD can achieve 0.064 higher accuracy than random source selection.

### 4.2. Neural Decoding Visualization and Cognitive Interpretation

It is also important to understand why a model performs well and which brain networks are particularly important, e.g. for advances in cognitive neuroscience and understanding neural disorder physiology. Exploring what information tends to emerge through domain adaptation will provide insights into how these methods work and what cognitive similarity is being leveraged through domain adaptation. Therefore, we carried out two studies to examine where the important voxels are in the brain, and how much the voxel sets in the target domain, source domain, and adapted models overlap.

#### 4.2.1. Model Overlapping Study I

We firstly studied the case of task 3 vs 6 as target and task 22 vs 9 as source, which showed the biggest improvement in classification accuracy. Figure 7 shows the voxels with the top 1% wight magnitude in the four models (target SVM, source SVM, TCA+SVM, and TCA+CDSVM) and occurring in clusters of at least 20 voxels. The target and source domain SVMs place their important voxels in completely different areas. Not only is there no overlap, but the supra-threshold voxels in each model are sampled from different lobes of the brain: the target domain SVM is associated primarily with supra-threshold voxels in the frontal lobes, while the source domain SVM is associated primarily with supra-threshold voxels in the occipital lobe and sensory motor cortex. The distribution of coefficients is so different between source and target models, but adaptation can be nevertheless very effective.

**Figure 7:**
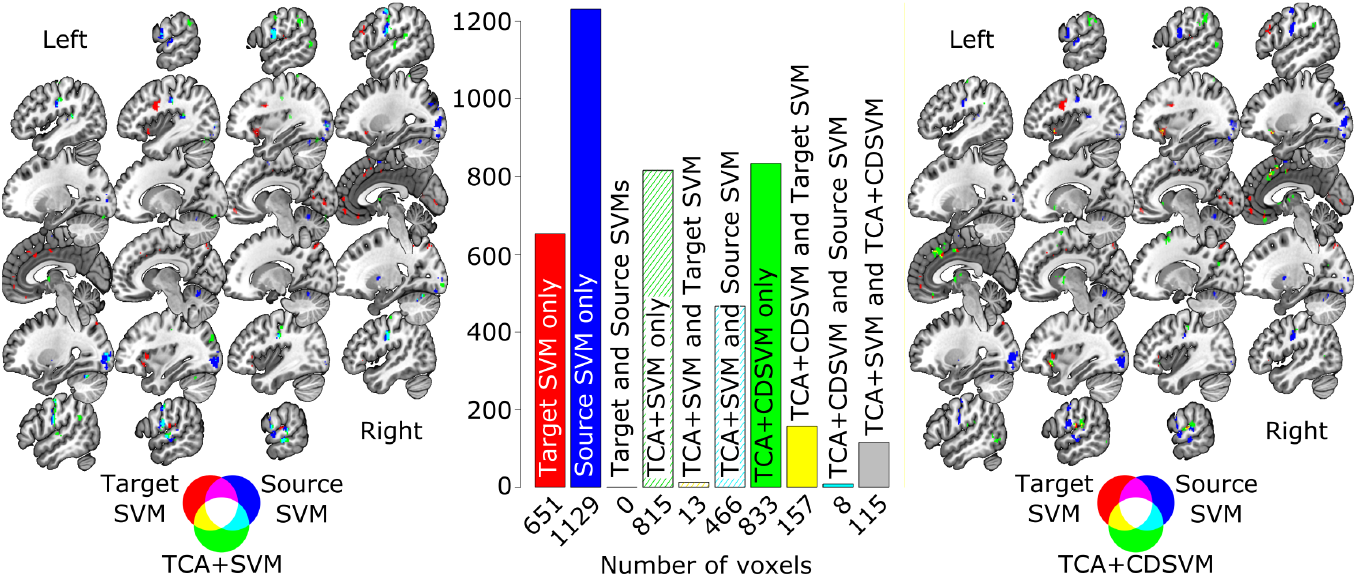
Visualization of the voxels with top 1% weight magnitude and occurring in clusters of at least 20 voxels in the four models: target SVM, source SVM, TCA+SVM, and TCA+CDSVM. Numbers of distinct and overlapped voxels identified by the models are shown in the middle bars.

We then overlaid the adapted models from TCA+SVM and TCA+CDSVM. TCA+SVM has substantial overlap with the source domain SVM, and nearly no overlap with the target domain SVM. This substantial overlap, however, is only about 1/3 of the supra-threshold voxels in the TCA+SVM model. The remaining 2/3 are completely distinct from either the target or source models, indicating that information from the source domain has revealed a different dimension along to which to dissociate the tasks in the target domain than was apparent in the target data in isolation.

This general pattern is echoed in the model adapted with TCA+CDSVM, except that in this case there is virtually no overlap with the source domain and there is instead modest overlap with the target domain. Again, the adapted model is largely associated with supra-threshold voxels that do not overlap with either the target or source SVMs. Thus, the adaptation procedure has provided additional insights into the classification problem, showing the exploited information to be more than the simple sum of information from the target and source domain models.

#### 4.2.2. Model Overlapping Study II

We further analyzed the overlapped important voxels (with top 1% weight magnitude) in the four aforementioned models for 142 different target-source pairs where TDSVM_Acc≤ 90% and TCA+CDSVM leading to at least 3% improvement. We examined the number of (overlapped) voxels for all 15 possible combinations of the four models, including individual models. We formed a 15-element vector for each target-source pair with 15 such numbers and then normalized it to unit length. Figure 8 depicts the normalized vectors as columns for all 142 target-source pairs, labeled with the corresponding model(s).

**Figure 8:**
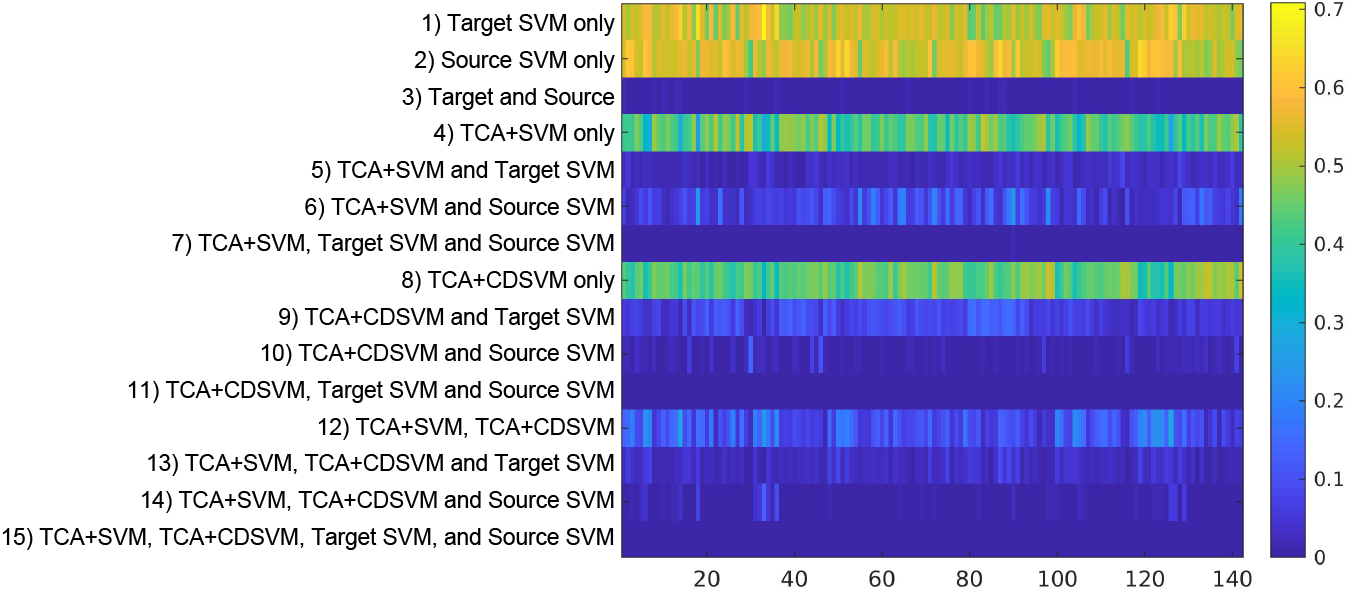
The overlapped important voxels for all 15 possible combinations (y-axis) of the four models: target SVM, source SVM, TCA+SVM, and TCA+CDSVM. The x-axis denotes 142 different target-source pairs where TDSVM_Acc≤ 90% and TCA+CDSVM leading to at least 3% improvement. The (overlapped) voxel numbers for the 15 model combinations form a 15-element vector for each target-source pair. This vector is normalized to unit length and visualized as a column. A larger value indicates a larger number of voxels.

There is a clear effect that each model identifies a fairly distinct set of voxels (rows 1, 2, 4, and 8), which are more than the overlapped voxels between nonadaptation and adaptation models (rows 5, 6, 9 and 10). On the other hand, by comparing the results shown in rows of 5, 6, 9 and 10, TCA+SVM overlaps with the source domain SVM much more than with the target domain SVM, and the opposite is true for TCA+CDSVM. This again confirmed that adaptation is exploiting information additional to the target and source domain models, and different adaptation schemes are exploiting different information.

### 4.3. Technical Challenges

Based on the experimental results, we can see two technical challenges in DawfMRI. One is *how to select a good source domain automatically (without exhaustive testing) to reduce or even avoid “negative transfer”*, as observed in our experiments. The plausible relationship between psychological similarity and adaptation effectiveness in Sec 4.1 can lead to a better than random solution. However, it is not the optimal selection of sources. The other challenge is *how to make use of multiple source domains to further improve the classification performance*. We can see such a need from the 2-CV results in Fig. 5, where domain adaptation is not effective when the number of training samples is very small. We need to carefully leverage the positive effects from each source domain while minimizing the potential negative impacts. This needs a smart, adaptive procedure to be introduced. We consider both challenges are important directions to explore in the future.

## 5 Conclusions

In this paper, we proposed a domain adaptation framework for whole-brain fMRI (DawfMRI). DawfMRI consists of three key steps: feature extraction, feature adaptation and classifier adaptation. We employed three state-of-the-art algorithms, ICA, TCA and CDSVM, for DawfMRI. We studied two nonadaptation algorithms and six adaptation algorithms on task-based whole-brain fMRI from seven OpenfMRI datasets. Results show that DawfMRI can significantly improve the classification performance for challenging binary classification tasks. We also observed “negative transfer” in the experiments, indicating that domain adaptation does not always give better performance and should be used with care. Furthermore, we discovered a plausible relationship between psychological similarity and adaptation effectiveness, and interpreted how the models provide additional insights. Finally, we pointed out two important research directions to pursue in future work.

## Acknowledgment

This work was supported by grants from the UK Engineering and Physical Sciences Research Council (EP/R014507/1) and Medical Research Council (MR/J004146/1), and the European Research Council (GAP: 670428 - BRAIN2 MIND_NEUROCOMP). The authors would like to thank the OpenfMRI (now OpenNeuro) project for providing the public fMRI datasets, and Professor Russell Poldrack and his team for providing help in reimplementing the OpenfMRI preprocessing pipeline.

## Footnotes

1 https://openneuro.org/

2 https://www.humanconnectome.org/

3 We used the data from the OpenfMRI project (https://openfmri.org/), now known as OpenNeuro. We will use the name OpenfMRI in the rest of this paper.

4 Data used in this paper are available at: https://legacy.openfmri.org/dataset/, also in the new BIDS format at OpenNeuro: https://openneuro.org/public/datasets

5 Only 19 subjects were involved in run 2 for this task.

6 https://github.com/poldrack/openfmri

7 http://www.cognitiveatlas.org/

